# A distinct adolescent profile for activity and dopamine release in the nucleus accumbens during Pavlovian conditioning

**DOI:** 10.64898/2026.05.21.726881

**Authors:** Ethan W. Herring, Madalyn Hafenbreidel, Eesha D. Patel, Sandford Zeng, Tulasi Syamala, Paul Kupelian, Mary M. Torregrossa, Sara E. Morrison

## Abstract

Across species, adolescence is a time of heightened reward sensitivity and enhanced impulsivity and risk-taking. In adults, these behavioral features are linked with a tendency to approach and interact with reward-associated cues – a behavior known as sign tracking – which is thought to reflect the transfer of incentive salience from reward to cue. Counterintuitively, adolescents are less likely to exhibit sign tracking, compared with adults, and more likely to exhibit goal tracking, or approach to the site of reward. To investigate a possible neural basis for this age difference, we recorded the activity of individual neurons in the nucleus accumbens (NAc) of male and female rats during Pavlovian conditioning in adolescence and adulthood. In a separate group, we used a fluorescent indicator (GRAB_DA_) to measure dopamine release at the same ages. We found that cue-evoked NAc activity increased over the course of training in adolescents and then further in adulthood. The majority of adolescents were goal trackers or intermediates, for whom reward-evoked activity peaked during adolescence and declines in adulthood, correlating with increased prevalence and intensity of sign tracking. Meanwhile, cue-evoked dopamine release was markedly higher in sign trackers than in goal trackers at all time points. These results suggest that the progression from adolescence to adulthood may be accompanied by changes in the engagement of the mesolimbic dopamine system and/or the responsivity of NAc neural signaling to dopamine, contributing to limited sensitivity to reward cues, coupled with heightened sensitivity to primary rewards, in adolescent animals.

**Significance Statement:** Adolescence is a time of enhanced reward sensitivity, impulsivity, and risk-taking, making adolescents vulnerable to drug use and other risky behaviors. In adults, attraction to reward-associated cues – which can be modeled in animals using a behavior called sign tracking – plays an important role in risky behaviors. Surprisingly, we find that adolescents exhibit less sign tracking compared with adults. Here, we investigate the neural circuits underlying this age difference by monitoring neural activity and dopamine release in the nucleus accumbens (NAc), a key brain area for reward-seeking behavior, in the same animals as adolescents and as adults. We find that the majority of adolescents show a reduced neural sensitivity to reward cues, but a heightened neural response to the reward itself.

## Introduction

Across species, adolescence is a time of heightened sensitivity to rewards (Friemel et al., 2010; Doremus-Fitzwater and Spear, 2016; Walker et al., 2017), along with enhanced impulsivity (Burton and Fletcher, 2012), exploratory behavior (Simon et al., 2013; Westbrook et al., 2018), and risk-taking (Doremus-Fitzwater et al., 2010; Zeng et al., 2023). This behavioral profile is thought to help adolescents acquire independence and learn the rules of their environment (Doremus-Fitzwater et al., 2010; Simon and Moghaddam, 2015). However, it likely contributes to adolescents’ unique vulnerability to the initiation of substance use (Chambers et al., 2003; Chen et al., 2009; Walker et al., 2017), along with other risky behaviors such as online gambling (Steinberg, 2008; Marquez-Ramos et al., 2023). In adults, maladaptive behaviors such as substance use are often influenced by reward-associated cues (Saunders and Robinson, 2013; Barrus et al., 2016) – stimuli that, over time, have acquired motivational value via their connection to rewarding outcomes, such as drugs, money, or food. Indeed, an individual’s sensitivity to reward-associated cues, or cue reactivity, is predictive of some measures of substance use and gambling propensity (Tomie et al., 2008; Flagel et al., 2009; Swintosky et al., 2021; Cherkasova et al., 2024).

Sign tracking is an easily quantified behavior that can be used to measure cue reactivity in humans and animals (Colaizzi et al., 2020). In a Pavlovian conditioning setting in which the cue (for rodents, typically the extension of a lever) is physically separate from the site of reward (e.g., a food magazine), some subjects will approach and interact with the cue – a behavior known as sign tracking (Hearst and Jenkins, 1974) – while others will approach the site of reward, a behavior known as goal tracking (Boakes, 1977). Sign tracking (ST) is thought to reflect the transfer of incentive salience from the reward to the cue (Flagel et al., 2009), and a propensity towards ST has been linked with various forms of impulsivity (Lovic et al., 2011), risk-taking (Swintosky et al., 2021), and measures of substance use and relapse (Tomie et al., 2008; Beckmann et al., 2011; Saunders and Robinson, 2013; Saunders et al., 2013; Tunstall and Kearns, 2015).

The role of reward cues in adolescents’ motivational processes is not well understood on a behavioral or neural level. Although some studies have found that adolescents have increased behavioral and neural responses to reward cues (Sturman et al., 2010; Burton et al., 2011; Sturman and Moghaddam, 2011), we and others have found that adolescent animals are markedly <u>less</u> likely to exhibit ST behavior (Anderson and Spear, 2011; Anderson et al., 2013; Rode et al., 2019; Vigorito et al., 2022) – and more likely to exhibit goal tracking (GT) – compared with adults. Only when adolescents are under substantial stress (e.g., social isolation and food restriction) do their levels of ST sometimes match or exceed those of adults (Anderson et al., 2013; DeAngeli et al., 2017). Studies also suggest that adolescents have a lesser tendency towards habit formation evoked by cued instrumental conditioning (Serlin and Torregrossa, 2015; Towner et al., 2020), and are less influenced by reward cues during risky decision-making in a gambling task (Zeng et al., 2023).

The acquisition of behaviors directed towards reward cues, including ST and other forms of Pavlovian conditioned approach, depends on the mesolimbic dopamine system (Cardinal and Everitt, 2004). Sign-tracking behavior, but not goal-tracking behavior, requires dopamine release in the nucleus accumbens (NAc) (Saunders and Robinson, 2012), and there is evidence that sign tracker vs. goal tracker individuals show distinct patterns of NAc dopamine release over the course of training (Flagel et al., 2011). Furthermore, we have shown that individual neurons in the NAc have activity patterns that differ between ST and GT subjects, perhaps reflecting differential modulation by dopamine (Gillis and Morrison, 2019; Duffer et al., 2023; Herring et al., 2025). Notably, the mesolimbic dopamine system undergoes profound changes during the transition from adolescence to adulthood (Doremus-Fitzwater and Spear, 2016; Reynolds and Flores, 2021), such as changes in the balance of D1 vs. D2 dopamine receptors in the NAc (Tarazi et al., 1999; Tarazi and Baldessarini, 2000), that could affect many aspects of NAc activity and related behavior. The connectivity of the NAc undergoes changes as well: for example, it receives increasing input from the medial prefrontal cortex (Brenhouse et al., 2008).

These findings led us to hypothesize that differences in NAc activity, possibly influenced by disparate patterns of dopamine release, might account for the differences between adults and adolescents in their behavioral response to rewards and reward-associated cues. Therefore, we recorded the activity of individual neurons in the NAc and, in a separate population, NAc dopamine release, during the acquisition and expression of ST and GT behavior in adolescent animals; then, we recorded again in the same subjects after they reached adulthood. We identified differences in NAc activity and dopamine that were connected with age and/or behavioral patterns as likely contributors to a unique adolescent neurobehavioral profile.

## Materials and Methods

All animal procedures were approved by the University of Pittsburgh’s Institutional Animal Care and Use Committee.

### Subjects and Timeline

Subjects for electrophysiology experiments were 13 Long-Evans rats (9 male, 4 female) bred in-house or obtained from Charles River Laboratories at age P21. They were housed on a 12 h reversed light/dark cycle (lights on at 7 P.M.), and all experiments took place during the dark phase. Animals bred in-house were weaned at age P21 and all subjects underwent surgery at age P23-25, after which they were single-housed to prevent damage to cranial implants. After surgery, rats were allowed to recover for at least 7 days before commencement of behavioral training. Single-unit recordings took place in the age range P33-45. Subjects were retrained for 2-3 days before recording in adulthood (adult retest), which took place at age 13-14 weeks. One animal (male) was excluded from adult retest due to a lost headcap; two others (both male) did not provide neural data at adult retest because they no longer had isolatable neurons. One animal (male) was excluded from analysis due to misplaced electrodes.

For certain analyses, we used a comparison group of adult-trained Long-Evans rats (11 males) from a previously published data set (Duffer et al., 2023). All of these subjects were obtained as young adults from Charles River Laboratories; they underwent surgery and training identical to adolescents except that all procedures took place in adulthood (age 3-4 months).

Subjects for photometry experiments were 18 Long-Evans rats (10 male, 8 female) obtained from Charles River at age P21. They were housed on a 12 hr light/dark cycle (lights on at 4 A.M.) and experiments took place during the dark phase. Animals underwent surgery at age P24-25 and were allowed to recover for at least 14 days before behavioral training to allow time for viral expression. Adolescent photometry recordings took place in the age range P38-46. Similar to electrophysiology experiments, subjects were retrained for 1-2 days before recording again in adulthood at age 13-14 weeks. The adult retest took place in a subset of subjects (n = 8; 4 males, 4 females). Three animals were excluded from analysis due to lack of viral expression (1 female; part of adult retest group) or misplaced expression/fiber optic (2 males; neither were part of adult retest group).

For both experiments, animals were placed on a restricted diet 2-3 days before behavioral training, consisting of 10 g/day of chow for adolescents and 14 g/day for adults. Rats were weighed regularly and provided with extra food if necessary to maintain at least 90% of expected body weight (as determined from standard growth charts for adolescents). Following the completion of training in adolescence, rats were given *ad libitum* food until 13-14 weeks of age, when they were returned to a restricted diet starting 3 days before retraining.

### Apparatus and Behavior

For electrophysiology experiments, all training and testing took place in the same operant chamber (Coulbourn Instruments) equipped with a house light, speaker for auditory cues, and a pellet dispenser connected to a food magazine recessed into the wall. The food magazine contained an infrared photobeam to detect magazine entries and exits. A retractable lever was installed on one side of the magazine with a cue light above it. Behavioral tasks were controlled by Coulbourn software (Graphic State 4.0).

Adolescent rats were initially given 2 days of magazine training, consisting of 50 noncontingent deliveries of a sucrose pellet (45 mg, BioServ) given at variable time intervals averaging 60s. During the second day of magazine training, rats were habituated to the headstage cable (Plexon) for neural recording. Subsequently, rats completed 7 or 8 d of the Pavlovian conditioned approach (PCA) task as we have previously described (Gillis and Morrison, 2019; Duffer et al., 2023). Briefly, the behavioral protocol started with illumination of the house light and consisted of 25 trials; intertrial intervals were selected from a truncated exponential distribution averaging 60s. Each trial began with an 8 s cue light illumination and lever extension, along with a tone at the beginning of the cue (1 s, 500-Hz intermittent tone). Immediately following cue cessation, a sugar pellet reward (40 mg, Bio-Serv) was delivered to the food magazine. No action was required for reward delivery.

For photometry experiments, the task and training were identical to the above, except training/testing took place in a different single operant box (Med Associates). Behavioral tasks were controlled by Med Associates software (MedPC).

### Surgical procedures

We employed standard aseptic surgical procedures. All animals were anesthetized using isoflurane (4% for induction, 1-2% for maintenance). Subjects used in electrophysiology experiments were treated with ketoprofen (5 mg/kg) and enrofloxacin (10 mg/kg) for 3 d post-surgery. Adolescent rats were implanted with an electrode bundle targeting the NAc core (coordinates in mm from bregma: AP, +1.0; ML, ± 1.2; DV, −6.5 from skull). Electrode bundles were constructed in house and consisted of 16 Teflon-insulated tungsten wires (A-M Systems) hand-cut to achieve an impedance of 90-110 KΩ.

Subjects used in photometry experiments were treated with carprofen (5 mg/kg) for 3 d post-surgery. Adolescent rats received an infusion of a viral vector encoding the fluorescent dopamine indicator GRAB_DA_ (AAV9-hSyn-GRAB_DA2m; Addgene) into the NAc core (coordinates in mm from bregma: AP, +1.0; ML, ± 1.2; DV, −6.2 from skull). A volume of 0.5 µL was infused at a rate of 0.1 µL per minute followed by a 5 min diffusion period. Subjects were then implanted with an optical fiber (2.5 mm ferrule, 400 µm core; Thorlabs) at the same coordinates. All photometry recordings took place at least 14 d following viral infusion.

### Electrophysiology

We recorded neural activity throughout all PCA task sessions using Plexon hardware and software. Rats were connected to a lightweight headstage that plugged into a commutator above the operant chamber to allow for free movement. Voltages were bandpass filtered between 220 Hz and 6 kHz, amplified 500x, and digitized at 40 kHz. Putative spikes were time-stamped and stored in segments of 1.4 ms. Units were hand-sorted (Offline Sorter; Plexon) using principal component analysis and visual inspection of waveform clusters. Units were analyzed only if they were >75 μV, had a signal-to-noise ratio of at least 2:1, and had fewer than 0.1% of interspike intervals <2 ms. Isolation of units was verified using autocorrelograms, as well as cross-correlograms for units recorded on the same electrode.

### Photometry

Photometry recordings were obtained from adolescents during the first training session (day 1), a middle training session (day 4 or 5), and the last training session (day 7 or 8), and again in adulthood (“adult retest”) after 2-3 days of retraining. We used a multi-wavelength fiber photometry system (Plexon) with a low-autofluorescence fiber optic patch cable (Doric; 400 µm core, 440 µm cladding, 0.37 NA). Rats were habituated to the patch cable during the second session of magazine training. Excitation wavelengths were 465 nm (GRAB_DA_) and 410 nm (isosbestic control) set to an intensity of 10-30 µW at the cable tip. Light was passed through the patch cable for 30 min prior to recording to minimize autofluorescence during recordings. Fluorescence data were collected at 30 frames per second using Plexon software.

### Histology

After completion of data collection, animals were deeply anesthetized with pentobarbital and transcardially perfused with saline followed by 10% formalin or 4% paraformaldehyde. Brains were post-fixed for 1 d and then transferred to 30% sucrose for at least 3 d before being cryostat-sectioned at 50 µm. Electrode tips were labeled by passing direct current through each electrode (75 µA for 10 s) prior to perfusion. Coronal NAc slices were stained with cresyl violet and electrode placements were confirmed with light microscopy.

Following photometry experiments, brain sections were imaged with an Olympus VS200 microscope at 10x magnification to confirm GRAB_DA_ expression and fiber placement.

### Analysis

Analyses were performed using custom-written programs in MATLAB (Mathworks). Significance was set at p ≤ 0.05 for all analyses unless otherwise specified.

#### Behavior

We quantified sign tracking and goal tracking using raw behavioral event counts (lever deflections and magazine entries) as well as a composite PCA index (Meyer et al., 2012). The PCA index is an average of three indices: the probability index is calculated as *P_lever_ -P_magazine_*, where *P* is probability of the indicated action; the bias index is calculated as (#lever press - #magazine entry)/ (#lever press + #magazine entry); and the latency index is calculated as (magazine latency – lever latency)/ (cue length). For trials with no behavioral response, the latency for that behavior was defined as the cue length (8 s).

#### Electrophysiology

Event-excited and event-inhibited neurons were identified as previously described (Gillis and Morrison, 2019; Duffer et al., 2023; Herring et al., 2025). Briefly, we defined a Poisson distribution approximating each neuron’s baseline firing rate in the 1 s before the event. Excited and inhibited neurons were identified by the presence of at least three consecutive 10 ms bins in which the firing rate exceeded the upper or was less than the lower 99.9% confidence interval of the baseline distribution, respectively. If both excitatory and inhibitory responses were identified within 500 ms after the event, we examined the mean *Z*-score over 200 ms, 500 ms, and 1 s post-event. If at least two consecutive time windows had positive average *Z*-scores, the neuron was categorized as event-excited; if at least two consecutive time windows had negative average *Z*-scores, the neuron was categorized as event-inhibited. Neurons with *Z*-scores that “flipped” from negative to positive to negative, or vice versa, over the three time windows had particularly complex responses and are shown as “uncategorized.”

Peristimulus time histograms (PSTHs) for individual neurons were calculated in 10 ms bins and are shown smoothed using a 5 bin moving average. Activity contributing to population PSTHs was calculated in 10 ms bins and Z-scored relative to baseline (1 s prior to event onset) before averaging; averaged activity is shown smoothed using a 5 bin moving average.

#### Photometry

Initial processing of photometry data was performed using GuPPy (Sherathiya et al., 2021). Briefly, fluorescence data were visualized and motion artifacts removed, then isosbestic data were fitted to the signal and subtracted to generate ΔF/F. Data were then Z-scored across the entire signal trace. Z-scored data were further processed using custom-written scripts in MATLAB (Mathworks) to visualize individual and average peri-event histograms and heat plots, peaks, and area under the curve (AUC) associated with behavioral and task events. Area under the curve was calculated over a 2 s window following an event, and peaks were derived from the same window.

After initial data processing, we determined that, in a number of subjects, signal was frequently lost during magazine entries associated with pellet consumption. (Signal during other magazine entries was minimally affected, which we verified via careful inspection of raw vs. processed signal and isosbestic traces.) Therefore, we opted to focus our analyses on GRAB_DA_ signal associated with cue presentation rather than with reward delivery/consumption.

## Results

In order to assess possible age-related changes in NAc activity during Pavlovian conditioning, we implanted adolescent rats (age P23-25; n=13) with microwire electrode bundles targeted to the nucleus accumbens (NAc) core. After recovery from surgery, rats were trained for 7 days on a Pavlovian conditioned approach (PCA) task that typically elicits sign-tracking and/or goal-tracking behavior (Gillis and Morrison, 2019; Duffer et al., 2023; Herring et al., 2025). Then, after a hiatus, the same rats were retrained on the PCA task for at least 2 days as adults (age 13-14 weeks). A timeline of experiments can be found in Figure 1A. Recording took place during all PCA task sessions; we focused our behavioral and neural analyses on Session 1, Session 7, and an adult retest (day 2 or 3 of retraining as adults).

**Figure 1.**
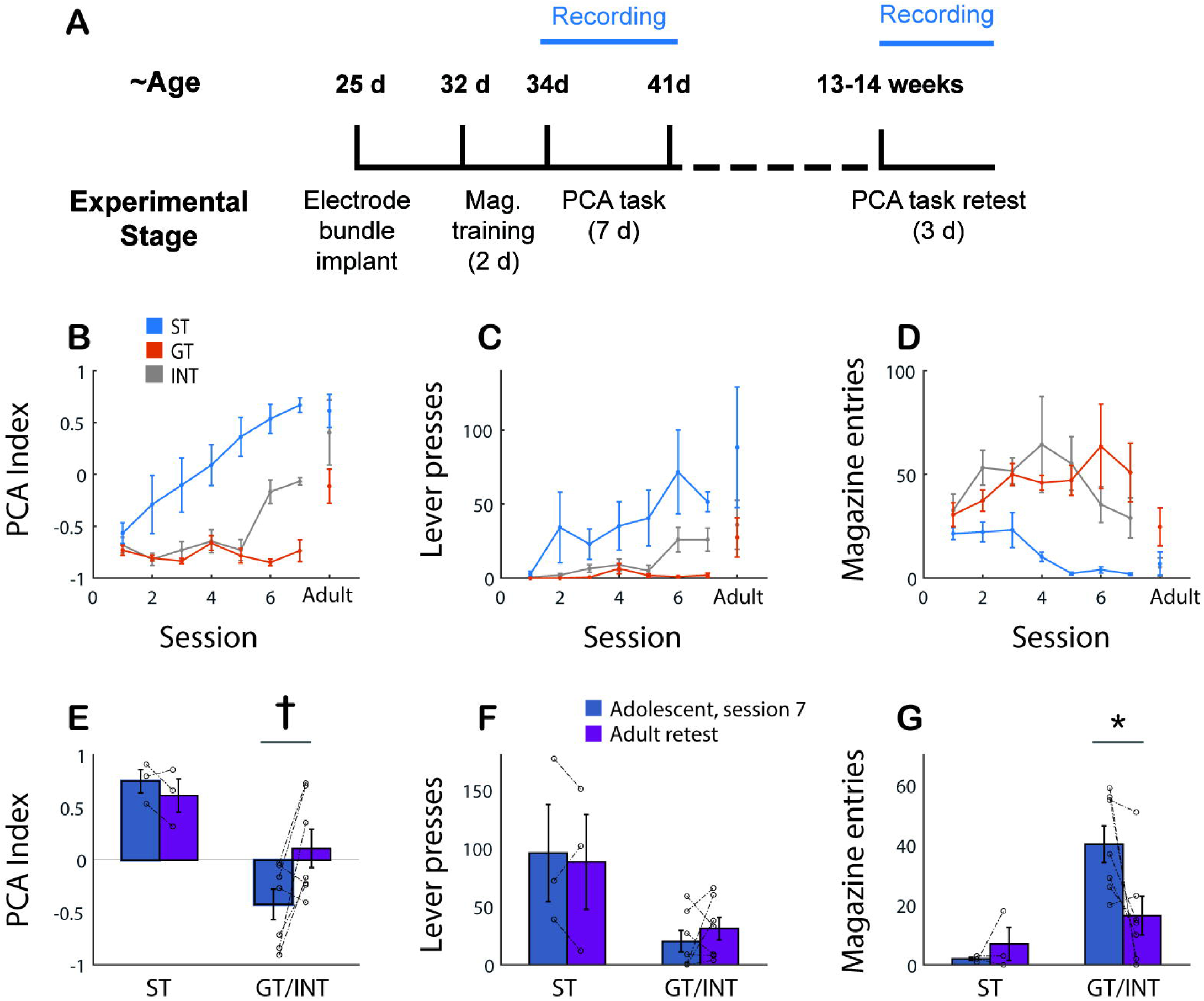
Adolescents frequently increase sign-tracking behavior and decrease goal-tracking behavior as they become adults. (**A**) Timeline of experiments. (**B-D**) Learning curve for PCA index (B), lever presses (C), and magazine entries (D) across 7 days of training in adolescence and at adult retest. Error bars, SEM. Blue indicates sign trackers, orange indicates goal trackers, and gray indicates intermediates as categorized on the last day of adolescent training. Some adolescent goal trackers became adult sign trackers. (**E-G**) PCA index (E), lever presses (F), and magazine entries (G) on the last day of training as adolescents (blue) and at adult retest (purple) for adolescent sign trackers (ST) and adolescent goal trackers / intermediates (GT/INT). Error bars, SEM. Asterisk, p = 0.03; dagger, p = 0.06, Wilcoxon signed rank test.

Sign tracking (ST), or approach/interaction with the cue, was represented by lever presses, while goal tracking (GT), or approach to the site of reward, was represented by entries into the food magazine during cue presentation. As we and others have done previously (Meyer et al., 2012), we quantified ST and GT behavior using a PCA index: a composite measure ranging from −1.0 to 1.0, where positive values indicate relatively stronger ST and negative values indicate relatively stronger GT. We categorized individual adolescents as sign trackers (ST), goal trackers (GT), or intermediate (INT) using their PCA index from Session 7: individuals with a PCA index > 0.25 were categorized as ST, individuals with a PCA index < −0.25 were categorized as GT, and all others were categorized as INT. Among the 10 subjects that were evaluated in both adolescence and adulthood, 3 were categorized as ST, 4 as GT, and 3 as INT. Three additional subjects (1 ST, 1 GT, 1 INT) were evaluated in adolescence but not as adults.

Behavioral trajectories for PCA index, along with sign tracking actions (lever presses) and goal tracking actions (magazine entries) over the course of training (7 days) and adult retest are shown in Figure 1B-D. We observed that adolescents categorized as GT or INT, but not the few categorized as ST, showed clear behavioral changes between the end of adolescent training and adult retest, including an increase in PCA index (Fig. 1B). To quantify this change, we combined the GT and INT groups (Fig. 1E-G), and found that they showed a trend towards an increase in PCA index (Fig. 1E; Z = 1.86, p = 0.06, Wilcoxon signed rank test). This was primarily driven by a significant decrease in goal-tracking behavior, i.e., magazine entries (Fig. 1G; Z = −2.20, p = 0.03). These results are consistent with previous findings, by ourselves and others, showing that adolescents, compared with separate adult populations, tend to exhibit less ST and more GT behavior (Doremus-Fitzwater and Spear, 2011; Anderson et al., 2013; Rode et al., 2019; Zeng et al., 2023).

### Electrophysiological recording

We obtained activity from 271 NAc cells in 11 adolescent subjects during the first session of training, 314 cells in 12 subjects during the last session, and 112 cells in 9 subjects at adult retest. A reconstruction of recording locations is shown in Fig. 2A. Visualization of activity from all cells (Fig. 2B-D) revealed that cue-evoked excitatory activity (in yellow) intensified over the course of training and from adolescence to adulthood. Reward-evoked excitatory activity also intensified, although this trend was less obvious, partially because inhibitory activity (dark blue) also became stronger. Patterns became more evident when we divided subjects into those who already exhibited robust ST behavior as adolescents (n=4) and those who were categorized as GT or INT. Both groups showed an increase in the proportion of cells that had excitatory responses to the cue over the course of the experiment. Among the GT/INT group (Fig. 2E), the ratio of cue-excited to cue-inhibited cells significantly increased from session 1 to session 7 (chi-square test: Χ^2^_(2,243)_ = 6.26, p = 0.04) and from session 7 to adult retest (Χ^2^_(2,216)_ = 6.00, p = 0.05). Among the ST group (Fig. 2F), the ratio was stable from session 1 to session 7 (Χ^2^_(2,306)_ = 2.21, p = 0.33), but increased from session 7 to adult retest (Χ^2^_(2,188)_ = 6.16, p = 0.05).

**Figure 2.**
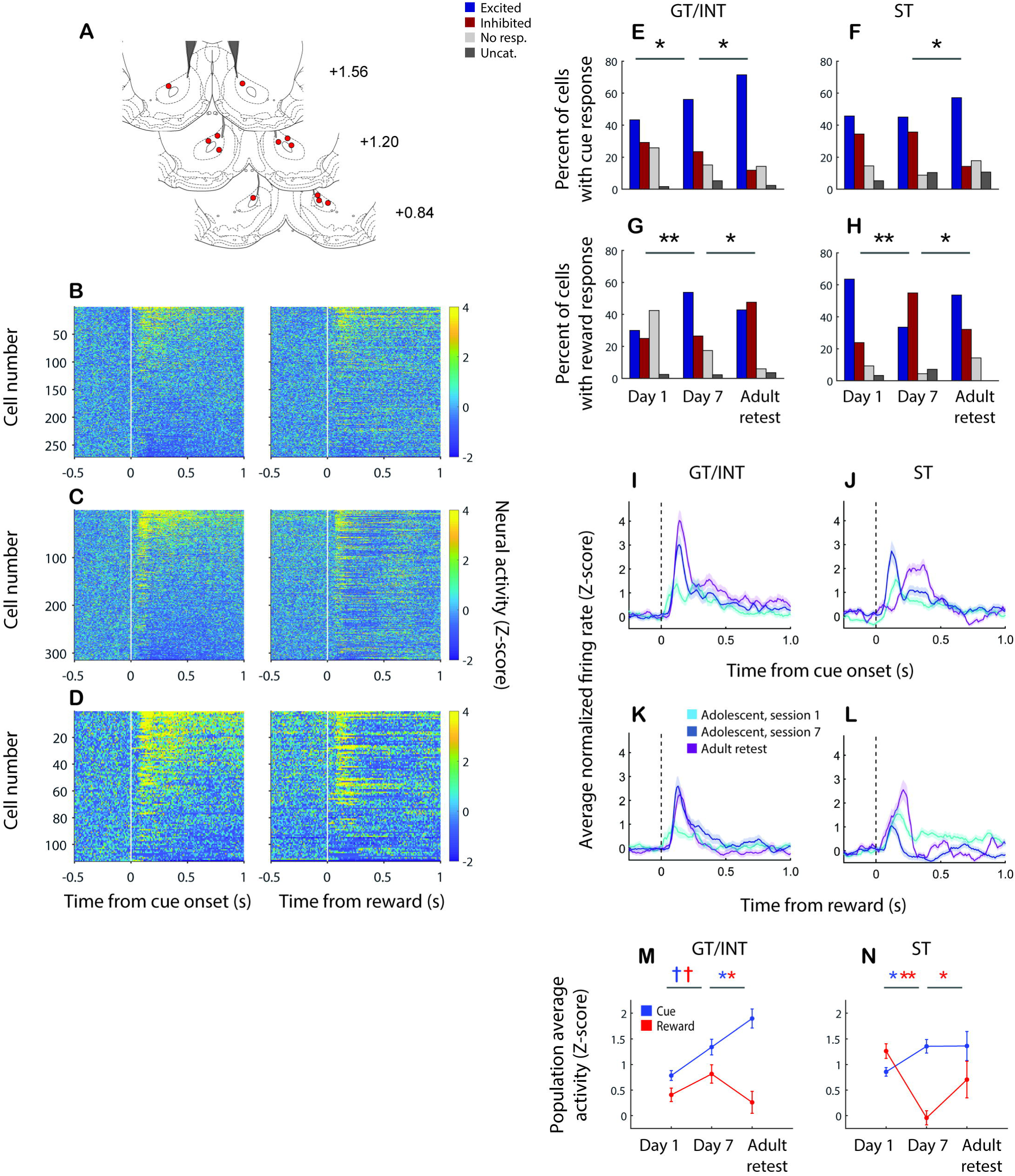
NAc excitatory cue responses increase from adolescence to adulthood, while reward responses peak in adolescence. (**A**) Coronal atlas images (Paxinos and Watson, 2007) showing the approximate location of electrodes in the NAc core. Numbers are distance in mm from bregma. (**B-D**) Average normalized activity of each neuron recorded on the first day (B) and last day (C) of training as adolescents and at adult retest (D) around the time of cue onset (left) and reward delivery (right). Activity is calculated in 10ms bins with no smoothing. Cells are sorted by cue response (average activity in the 1s following cue onset). (**E-H**) Percent of cells identified as having an excitatory response (orange) or inhibitory response (blue) to the cue (E,F) or reward (G,H) among subjects categorized as GT/INT (E,G) or ST (F,H). Light gray, no significant response; dark gray, uncategorized. Asterisk, p ≤ 0.05; double asterisk, p ≤ 0.001, chi square test. (**I-L**) Population average normalized activity of cue-excited neurons at cue onset (I,J) and reward delivery (K,L) among subjects categorized as GT/INT (I,K) or ST (J,L). First day of adolescent training, light blue; last day of adolescent training, dark blue; adult retest, purple. Shading indicates SEM. (**M,N**) Population average activity in the 1s following cue onset (blue) or reward delivery (red) at each of the three timepoints for GT/INT (M) and ST (N) subjects. Error bars, SEM. Dagger, p ≤ 0.1; asterisk, p ≤ 0.05; double asterisk, p ≤ 0.001, Wilcoxon rank sum test. Blue dagger/asterisks indicate changes in cue response, red dagger/asterisks indicate changes in reward response.

Reward responses were more sharply divergent between the two groups. Among GT/INT subjects (Fig. 2G), a large number of cells had little or no response to reward early in training, but the proportion of cells that showed excitatory responses to reward increased over the course of adolescent training (chi-square test: Χ^2^_(2,246)_ = 21.90, p < 0.001), peaking at session 7. Later, at adult retest, GT/INT animals exhibited fewer reward-excited cells and an increase in reward- evoked inhibitory responses (Χ^2^_(2,210)_= 13.06, p = 0.001). ST subjects, in contrast (Fig. 2H), had more cells exhibiting reward-evoked excitations on the first day of training, followed by a decrease in excitations and a large increase in inhibitions by the end of adolescent training (Χ^2^_(2,315)_ = 38.08, p < 0.001); then, by adult retest, the proportions of excitations and inhibitions were reversed (Χ^2^_(2,197)_= 8.67, p = 0.01).

This general pattern was evident even when we restricted our analyses to neurons with cue-evoked excitatory responses, as we have in prior studies (Gillis and Morrison, 2019; Duffer et al., 2023). We have previously shown that the population activity of cue-excited neurons (which comprises about half of recorded NAc neurons in adults) differs between ST and GT animals (Gillis and Morrison, 2019). In adolescents, we likewise saw a divergent response profile (Fig. 2I-N). Among both GT/INT and ST groups, the population cue-evoked excitatory response increased markedly from early to late training in adolescence (Fig. 2I,J). On a per cell basis, the average firing rate over the 1s following cue onset was significantly increased in the ST group (Fig. 2N; Z = 2.32, p = 0.02, Wilcoxon rank sum test) and trended towards an increase in the GT/INT group (Fig. 2M; Z = 1.64, p = 0.10). Among GT/INT subjects, cue-evoked excitation increased further between adolescence and adulthood (Fig. 2I); in contrast, among ST subjects, cue-evoked responses stayed level in magnitude from adolescence to adulthood (Fig. 2J), even though cue-evoked excitations occurred in a larger proportion of cells (Fig. 2F). Likewise, on a per cell basis, the average cue-evoked firing rate significantly increased in the GT/INT group (Fig. 2M; Z = 2.28, p = 0.02), but not in the ST group (Fig. 2N; Z = 0.33, p = 0.74).

Among the population of cue-excited cells, GT/INT subjects also showed an increase in reward-evoked excitation over the course of adolescent training, which then diminished in adulthood (Fig. 2K); specifically, their adult reward responses reached a similar peak to those of adolescents, but were not sustained as strongly. On a per cell basis, the average firing rate in the 1s following reward delivery trended towards an increase over the course of adolescent training (Fig. 2M; Z = 1.74, p = 0.08), then significantly decreased in adulthood (Z = −2.66, p = 0.008). In ST subjects, on the other hand, reward-evoked responses markedly diminished between the first day and last day of adolescent training, then rebounded in adulthood (Fig. 2L). On a per cell basis, the average reward-evoked firing rate significantly decreased over the course of adolescent training (Fig. 2N; Z = −6.20, p < 0.001) and increased in adulthood (Z = 1.98, p = 0.05).

Taking these results together, when we consider the adolescents most typical of the wider population – i.e., the GT/INT group (Doremus-Fitzwater and Spear, 2011; Anderson et al., 2013; Rode et al., 2019) – we see that this group’s cue-evoked NAc activity increased between adolescence and adulthood; on the other hand, reward-evoked activity peaked in adolescence, then decreased in adulthood. These activity patterns were similar to those we have previously observed in adult ST subjects during learning – particularly the reduction in reward-evoked excitation (Gillis and Morrison, 2019) – that are in turn similar to patterns of cue- and reward-evoked dopamine release in adult ST animals (Flagel et al., 2011). On the other hand, in the group that developed strong ST behavior as adolescents, the same pattern – increased excitation to the cue and decreased excitation to reward – occurred <u>within</u> adolescence.

We next investigated whether changes in event-related NAc activity were related to changes in behavior over the course of development. Indeed, among individual subjects, change in PCA index between sessions showed a significant negative correlation with average change in reward-evoked neural activity (Fig. 3B; r^2^ = 0.19, p = 0.05). There was also a positive relationship with cue-evoked activity (Fig. 3A), although it did not rise to the level of significance (r^2^ = 0.12, p = 0.14). Finally, there was a robust positive relationship between increasing PCA index and an increasing <u>difference</u> between cue-evoked and reward-evoked neural responses (Fig. 3C; r^2^ = 0.33, p = 0.008), implying that increases in sign tracking, relative to goal tracking, were associated with a shift in NAc excitation from reward to cue.

**Figure 3.**
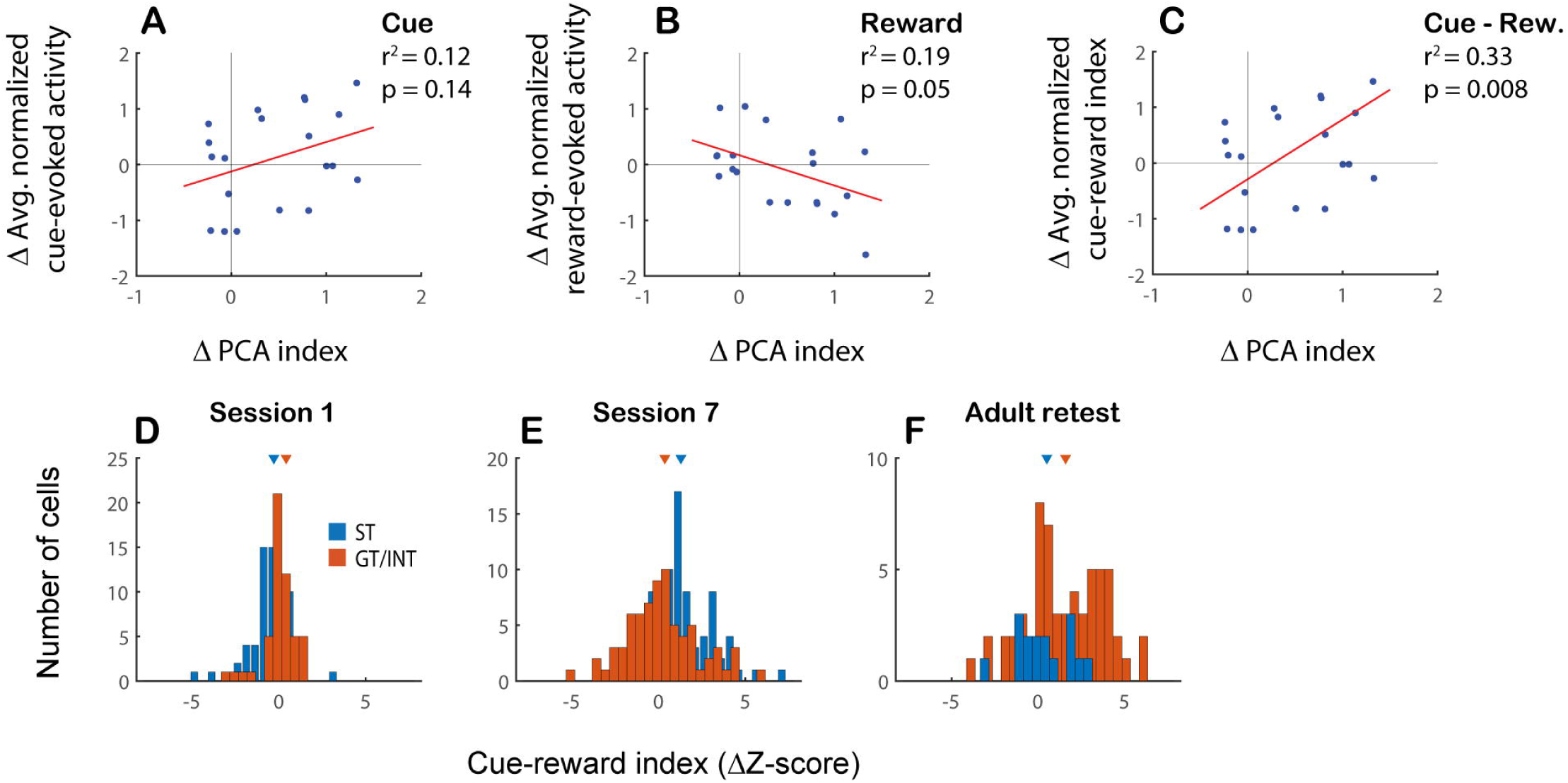
Increased sign tracking is associated with increased cue-evoked and reduced reward-evoked activity in the NAc. (**A-C**) Change in PCA index across sessions plotted against concurrent change in average neural activity evoked by cue onset (A) or reward delivery (B) or cue-reward index, defined as the difference between cue-evoked and reward-evoked activity (C). Each subject contributes a maximum of two data points to each plot (session 1 to session 7; session 7 to adult retest). Regression lines in red. (**D-F**) Distribution of cue-reward index during session 1 (D), session 7 (E), and adult retest (F). GT/INT group in orange; ST group in blue. Orange arrowhead, median of GT/INT distribution; blue arrowhead, median of ST distribution. The distribution shifts in a positive direction for only the ST group between session 1 and session 7 (p < 0.001, Wilcoxon rank sum test), and for only the GT group between session 7 and adult retest (p = 0.003).

Similarly, across the population of cue-excited neurons, differences between cue-evoked and reward-evoked responses in the same cell (cue-reward index) exhibited different distributions across stages of learning and among ST vs. GT/INT subjects. The distribution of cue-reward index was initially significantly shifted from zero in both groups (Fig. 3D; Session 1: ST: Z = - 2.82, p = 0.005; GT: Z = 4.16, p < 0.001, Wilcoxon signed rank test), but in opposite directions: ST subjects started out with greater reward responses (relative to cue), while GT/INT subjects started out with greater cue responses (relative to reward). But by the last day of training as adolescents (Fig. 3E), the cue-reward index for ST subjects was significantly shifted from zero in the positive direction (Z = 5.67, p < 0.001), while the index for GT/INT subjects was just barely shifted from zero (Z = 1.93, p = 0.054). Then, upon retesting as adults (Fig. 3F), the index for ST subjects was no longer significantly different from zero (Z = 1.45, p = 0.15), but was shifted in the positive direction for the GT/INT group (Z = 4.69, p < 0.001) The distributions for ST vs. GT/INT subjects were also significantly different from each other in adolescence (Session 1, Z = 4.72, p < 0.001; Session 7, Z = 2.89, p = 0.003, Wilcoxon rank sum test) and not in adulthood (Z = 1.69, p = 0.09).

Across sessions, the shift in cue-reward index from adolescent Session 1 to Session 7 was highly significant in ST subjects (Z = 6.59, p < 0.001, Wilcoxon rank sum test) but not in GT subjects (Z = 0.08, p = 0.94). Conversely, the shift in cue-reward index from Session 7 to Adult Retest was not significant in ST subjects (Z = 1.43, p = 0.15) but significantly shifted in a positive direction in GT/INT subjects (Z = 2.95, p = 0.003). Thus, NAc neural activity showed a strong transfer from reward to cue during adolescent training among ST subjects; however, among GT/INT subjects, the relationship between cue- and reward-evoked responses did not appreciably change during adolescence. Instead, among GT/INT subjects, activity transferred from reward to cue in adulthood.

Finally, we noted that adolescents showed remarkably muted NAc activity in response to events, including cue and reward delivery, at the beginning of training, despite our previous observations that NAc neurons are often excited by novel stimuli (Gillis and Morrison, 2019; Duffer et al., 2023). Therefore, we used a preexisting set of recordings from adult male subjects (Duffer et al., 2023) (n = 11) to compare the evolution of signals in animals trained in adolescence with those trained in adulthood. While NAc neurons recorded in adults (n = 69 cue-excited cells) showed robust excitatory responses to both cue and reward during the first session of training, adolescents had far weaker responses, even among those cells algorithmically identified as cue-excited (n = 93; Fig. 4A,B). This might contribute to the significantly lower level of behavior seen in adolescents, compared with adults, on the first day of training (Fig. 4C; Wilcoxon rank sum, Z = 2.62, p = 0.009 for magazine entries).

**Figure 4.**
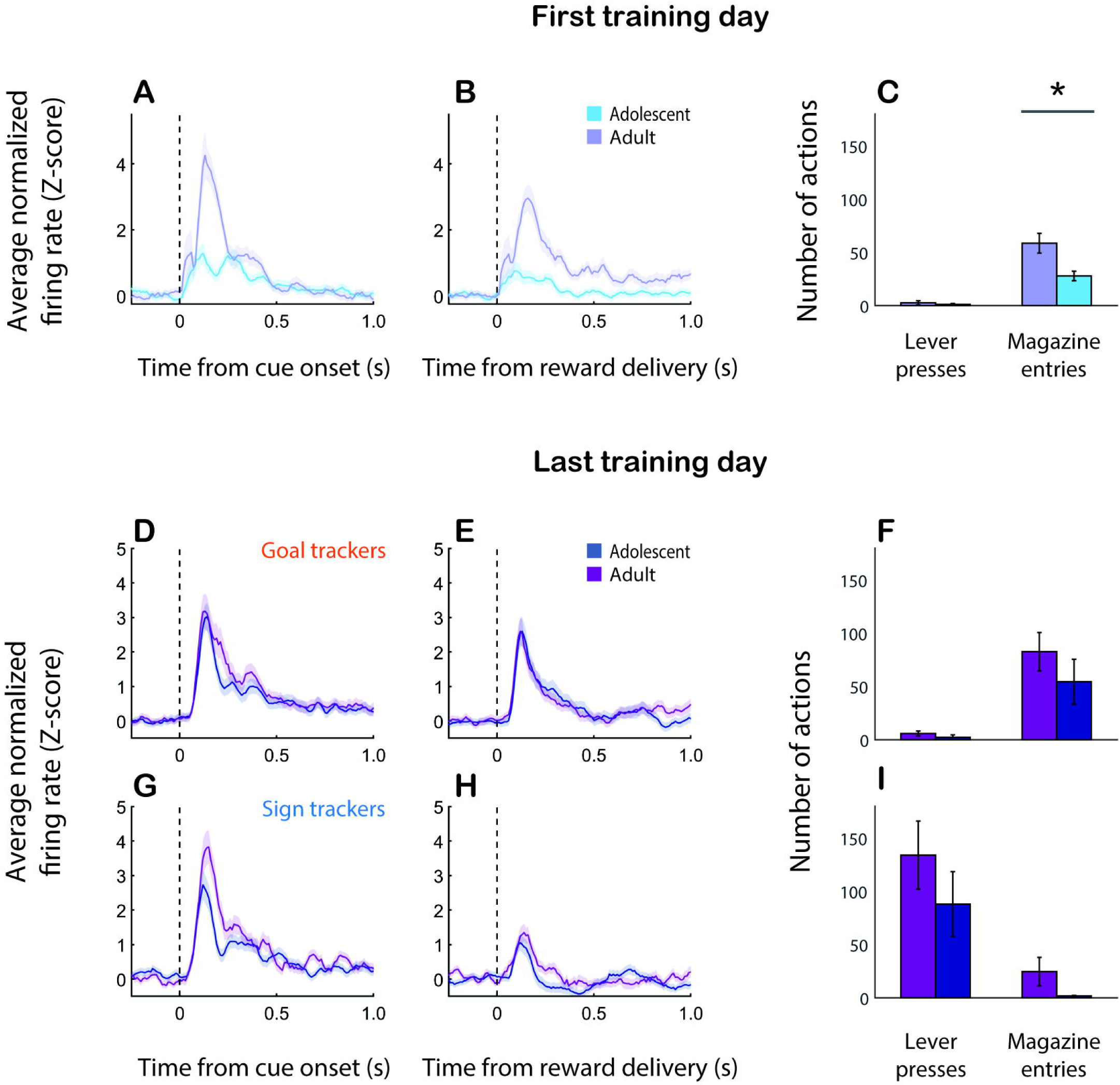
Comparison with adult-trained subjects. (A-C) Population average normalized activity in response to the cue. (A) and reward (B), and average behavior (C), on the first day of training for adolescents (blue) and a separate population of adult subjects (purple). Shading and error bars both indicate SEM. Asterisk, p = 0.009, Wilcoxon rank sum test. (**D-I**) Population average normalized activity in response to the cue (D,G) and reward (E,H), and average behavior (F,I) on the last day of training among adolescents (blue) categorized as sign trackers (D-F) or goal trackers (G-I) and a separate population of adults categorized as sign trackers or goal trackers. INT subjects not included. Shading and error bars both indicate SEM.

In contrast, by the last day of training (Session 7), when divided into ST and GT subjects, there were only small differences in cue- and reward-evoked responses between the adult-trained and adolescent-trained groups (Fig. 4D,E,G,H) – most notably, slightly stronger responses to cues in adults of both groups, on average. This is consistent with the non-significant differences in behavior between adult vs. adolescent GT subjects (Fig. 4F; Z = 1.04, p = 0.30 for magazine entries) and adult vs. adolescent ST subjects (Fig. 4I; Z = 1.24, p = 0.23 for lever presses). Taken together, these data account for our finding that adolescents show robust increases in cue-evoked excitation over the course of learning (Fig. 2I,J,M,N) even though we have previously reported only slight increases in cue responding in adults (Gillis and Morrison, 2019), along with strong decreases in reward-evoked excitation among ST animals.

### Fiber photometry

A number of studies have shown that dopamine release in the NAc core is essential for the acquisition of sign tracking, but not goal tracking (Saunders and Robinson, 2012; Iglesias et al., 2023; Herring et al., 2025), and we and others have observed that the cue- and reward-related activity of individual NAc neurons is modulated by dopamine (du Hoffmann and Nicola, 2014; Herring et al., 2025). Therefore, we hypothesized that patterns of NAc dopamine release might be related to the changes in ST and GT behavior that occur during adolescent maturation. To test this hypothesis, we expressed the fluorescent dopamine sensor GRAB_DA_ unilaterally in the NAc core of adolescent rats (n = 18), allowing at least 2 weeks for viral expression. Then, using fiber photometry, we measured GRAB_DA_ fluorescence at three timepoints – early, mid, and late training sessions – during the same PCA task used in electrophysiology experiments. Finally, after a hiatus, we retrained a subset of subjects (n = 8) for 2-3 days as adults (age 13-14 weeks) before recording GRAB_DA_ signal during an adult retest.

The behavior of subjects that contributed photometry data (n = 15) is shown in Figure 5A-C. As in the previous experiment, we categorized subjects as ST (n = 4), GT (n = 5), or INT (n = 6) using their PCA index on the last day of adolescent training. Similar to earlier behavioral results, we noted that individuals categorized as ST in adolescence generally retained their ST behavior in adulthood. GT and INT adolescent subjects, on the other hand, often (but not always) increased their ST behavior and/or decreased their GT behavior in adulthood. This cohort of GT/INT subjects showed a more modest shift in behavior from adolescence to adulthood, on average, compared to subjects that participated in electrophysiology experiments (Fig. 1D-F); this was driven by several GT subjects that remained GT in adulthood. By chance, only one subject categorized as INT was retested in adulthood, at which time the subject had developed more robust sign-tracking behavior.

**Figure 5.**
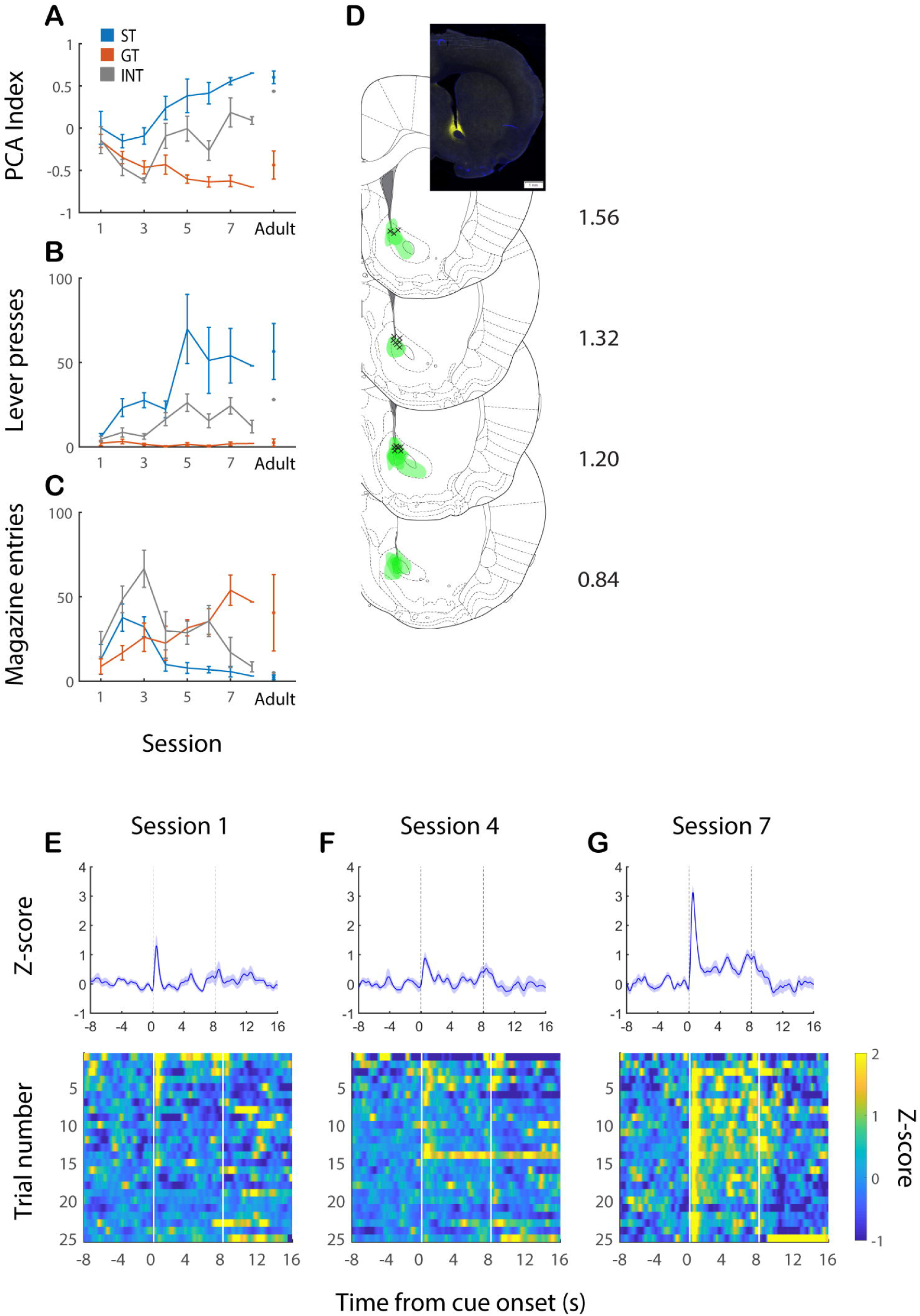
Behavior, histology, and representative results of GRAB_DA_ photometry. (A-C) Behavior, including PCA index. (A), total lever presses (B), and total magazine entries (C) during 7 sessions of adolescent training and at adult retest. Blue indicates sign trackers, orange indicates goal trackers, and gray indicates intermediates categorized on the last day of adolescent training. Error bars, SEM. Not all subjects are included in all data points; only one INT subject was present for adult retest. (**D**) Coronal atlas images (Paxinos and Watson, 2007) showing approximate spread of GRAB_DA_ construct expression (green) and fiber tip locations (black markers). Inset, representative example of GRAB_DA_ expression (green) in NAc core (blue is DAPI). Scale bar, 1mm. (**E-G**) Representative example GRAB_DA_ recordings in an adolescent on the first day (E), midpoint (F), and last day (G) of training. This subject was categorized as an intermediate, exhibiting both ST and GT behavior during the last day of training. Upper plots, average Z-scored GRAB_DA_ signal; shading indicates SEM. Lower plots, trial-by-trial heatmaps of Z-scored signal. Cue onset is at 0 s and cue offset / reward delivery are at 8 s.

GRAB_DA_ expression and fiber tip locations, assessed either in adolescence (n = 7) or after adult retest (n = 8), were concentrated in the dorsal and dorsomedial NAc core. The extent of expression was similar in adolescents and adults, so results were combined for display (Fig. 5D). Recordings of GRAB_DA_ fluorescence from a representative adolescent subject are shown in Figure 5E-G. This subject was categorized as an INT, exhibiting a mix of ST and GT behavior during session 7; lower levels of both behaviors were present during sessions 1 and 4. While a small phasic dopamine response to the cue was present early in training (Fig. 5E), it intensified by late training (Fig. 5G), with dopamine remaining somewhat elevated throughout the cue period. Meanwhile, only minimal responses to cue offset and reward delivery were present. Although the example recordings shown are relatively free of artifacts, we were unable to assess GRAB_DA_ signal in response to reward delivery across the wider subject population due to signal loss during magazine entries. Therefore, our analyses were primarily focused on GRAB_DA_ signal in response to cue presentation.

The pattern of cue-evoked dopamine seen in the example recordings (Fig. 5E-G) was typical of INT subjects, as demonstrated by the population average GRAB_DA_ signal (Fig. 6A-D) and cue-related AUC and peak signal (Fig. 6E,F). In contrast, subjects categorized as ST in adolescence showed an elevated phasic dopamine response to the cue even early in training (Fig. 6A), which was maintained or slightly elevated over the course of adolescent training. Consistent with their lack of behavioral changes as they reached adulthood, ST subjects’ cue-evoked dopamine release remained stable or even decreased slightly (Fig. 6D-F). Meanwhile, subjects categorized as GT in adolescence showed only a small GRAB_DA_ response to the cue throughout training, including in adulthood. Accordingly, there was a significant main effect of adolescent behavioral phenotype (ST, GT, or INT) on cue-related AUC (Fig. 6E; F_(2,51)_ = 4.99, p = 0.01) and peak (Fig. 6F; F_(2,51)_ = 4.05, p = 0.03). In pairwise comparisons, the ST group exhibited a significantly higher AUC (Z = 2.02, p = 0.04, Wilcoxon rank sum test) and a trend towards a higher peak (Z = 1.89, p = 0.06) on the first day of training compared with all other subjects. In contrast, by the last day of adolescent training, the GT group had a significantly lower AUC (Z = 2.00, p = 0.046) and a trend towards a lower peak (Z = 1.87, p = 0.06) compared with all other subjects.

**Figure 6.**
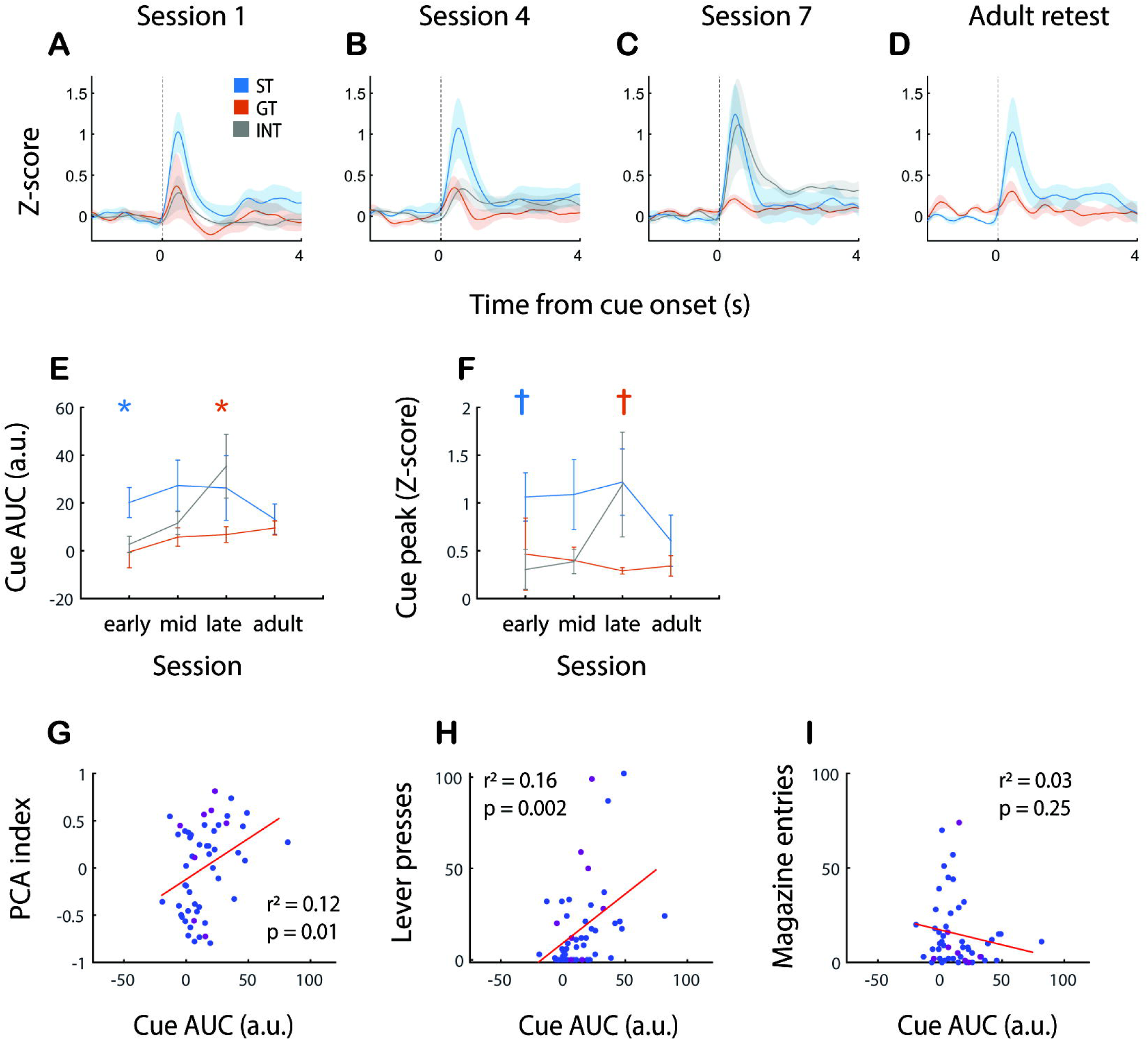
Cue-evoked dopamine is associated with sign-tracking behavior across age groups. (**A-D**) Average Z-scored GRAB_DA_ cue-evoked signal for subjects categorized in adolescence as sign trackers (blue), goal trackers (orange), or intermediates (gray) on the first day (A), midpoint (B), and last day (C) of adolescent training and at adult retest (D). Shading, SEM. (**E,F**) Average GRAB_DA_ signal area under the curve (E) and peak GRAB_DA_ signal (F) over the three adolescent test days and adult retest for sign trackers (blue), goal trackers (orange), and intermediates (gray). Blue symbols indicate ST group is different from combined GT/INT; orange symbols indicate GT group is different from combined ST/INT. Blue asterisk, p = 0.04; orange asterisk, p = 0.05; blue and orange daggers, p = 0.06. Error bars, SEM. (**G-I**) Scatter plots with regression lines (red) showing a significant correlation between the GRAB_DA_ AUC and PCA index (G) or total lever presses (H) but not magazine entries (I). Blue indicates adolescent time points; purple indicates adult time points.

Consistent with these findings, subjects’ cue-evoked GRAB_DA_ signal, averaged over each session, was significantly correlated with measures of cue-related behavior in that session, including PCA index (Fig. 6G; r^2^ = 0.12, p = 0.01) and total lever presses (Fig. 6H; r^2^ = 0.16, p = 0.002). Notably, however, it was not significantly correlated with total magazine entries (Fig. 6I; r^2^ = 0.03, p = 0.25). Thus, cue-evoked dopamine release was specifically related to sign tracking, rather than goal tracking, behavioral responses.

## Discussion

Adolescents, across species, are uniquely sensitive to rewards and relatively insensitive to aversive outcomes (Friemel et al., 2010; Doremus-Fitzwater and Spear, 2016; Palminteri et al., 2016). However, increasing evidence suggests that adolescents are less likely than adults to transfer motivational value from rewards to reward-predictive cues (Anderson et al., 2013; Rode et al., 2019; Zeng et al., 2023). In the current study, we investigated a possible neural basis of divergent behavioral responding to reward-associated cues in adolescent vs. adult animals. We found that many individual subjects increased their interaction with cues (sign tracking) and decreased their interaction with the site of reward (goal tracking) as they matured from adolescents into adults. This was accompanied by a robust increase in cue-evoked neural activity in the NAc, as well as a decrease in reward-evoked activity; moreover, the magnitude of these neural changes predicted the extent of behavioral change. Meanwhile, NAc cue-evoked dopamine release specifically scaled with the magnitude of ST behavior, suggesting that it is likely to promote the increase in incentive salience-driven behavior across developmental stages.

### Adolescent and adult behavior in response to reward-associated cues

Sign tracking and goal tracking are often conceptualized as the behavioral outputs of two different (but not mutually exclusive) learning processes (Clark et al., 2012; Huys et al., 2014; Lesaint et al., 2014). Goal tracking uses the predictive or informational content of the cue to guide behavior, but goal trackers do not assign the cue motivational value of its own. Sign tracking, on the other hand, is the result of a transfer of incentive salience from the reward to the cue itself, resulting in approach and interaction with the cue – in this case, an extended lever. This idea is supported by the finding that the cue acts as a potent conditioned reinforcer in sign trackers, but not goal trackers (Robinson and Flagel, 2009; Flagel et al., 2011; Meyer et al., 2012).

In adult animals, a propensity towards sign tracking has been linked with impulsive action (Lovic et al., 2011), risk-taking (Swintosky et al., 2021), novelty-seeking (Flagel et al., 2009), and drug-seeking (Flagel et al., 2009; Saunders and Robinson, 2013; Tunstall and Kearns, 2015). Many of these behavioral characteristics are similar to those of adolescents, who also show enhanced impulsivity (Andrzejewski et al., 2011; Burton and Fletcher, 2012), risk-taking (Steinberg, 2008; Westbrook et al., 2018; Zeng et al., 2023), and novelty- or sensation-seeking (Simon et al., 2013; Steinberg et al., 2018; Westbrook et al., 2018) compared with adults in both humans and animal models. However, counterintuitively, a number of studies have shown that adolescent animals are actually <u>less</u> likely to exhibit sign-tracking behavior than adults, and more likely to exhibit goal tracking (Anderson and Spear, 2011; Anderson et al., 2013; Rode et al., 2019). Moreover, we have previously shown that adolescent goal-tracking behavior is genuinely goal-directed – i.e., not a covert form of sign tracking – as defined by its sensitivity to reward devaluation (Rode et al., 2019). Indeed, in both adolescents and adults, goal tracking is far more sensitive than sign tracking to manipulations of reward value or of cue-reward contingency, such as extinction (Ahrens et al., 2015; Morrison et al., 2015; Patitucci et al., 2016; Gillis and Morrison, 2019).

A smaller propensity towards sign tracking implies that, compared with adults, adolescents are less likely to transfer incentive salience from a reward to a reward-predictive cue. This may be related to the observation that adolescents are less likely to develop habitual behavior following either cued or uncued instrumental conditioning (Serlin and Torregrossa, 2015; Towner et al., 2020). Both of these findings support the idea that adolescent behavior is biased towards flexibility and exploration (McCormick and Telzer, 2017; Baker et al., 2025), rather than exploitation (Harms et al., 2024), even at the expense of consistency, automaticity, and/or computational efficiency.

There is evidence that elements of the brain circuitry underlying sign tracking and other inflexible forms of behavior such as habit – in which the mesolimbic dopamine system is heavily implicated (Huys et al., 2014; Lerner, 2020) – are not fully mature in adolescents (Doremus-Fitzwater et al., 2010). It is also possible that, during adolescence, this circuitry may be more stringently gated by interoceptive or environmental conditions such as hunger or stress (Anderson et al., 2013; Hinton et al., 2019). In either case, we would expect the system for attributing incentive salience to cues to “come online” and dominate behavior more fully as animals mature from adolescents into adults. The current study provides support for this idea: individual animals often (but not always) increase sign tracking and decrease goal tracking from adolescence to adulthood. This results in many goal tracker and intermediate adolescents emerging as sign tracker adults; in contrast, adolescent sign trackers virtually never become goal trackers as adults.

### Neural substrates of cue responding in adults and adolescents

The NAc is essential for the acquisition and expression of taxic approach towards reward-associated cues (Nicola, 2010; Morrison et al., 2017). We and others have shown that the responses of individual neurons in the NAc core – in particular, cue-evoked excitations – scale with the probability and vigor (e.g., latency and speed) of behavioral responses (McGinty et al., 2013; Morrison et al., 2017). In adults, distinctions between sign trackers and goal trackers mainly emerge in reward-evoked activity: sign trackers show a marked decrease in reward-related excitation over the course of training that is not seen in goal trackers (Gillis and Morrison, 2019). This might reflect differential modulation by dopamine release: sign trackers reportedly show an increase in cue-evoked dopamine and a decrease in reward-evoked dopamine, resembling a reward prediction error (RPE) signal, whereas goal trackers do not (Flagel et al., 2011).

Here we report that adolescent sign trackers, like adult sign trackers, have an RPE-like decrease in NAc reward-evoked responses over the course of training. At the same time, they show an increase in cue-evoked responding that is more pronounced than the one we have seen in adults (Gillis and Morrison, 2019). Adolescent goal-tracker and intermediate subjects, on the other hand, show robust increases in both cue-evoked and reward-evoked activity over the course of training. Then, when retested in adulthood, they exhibit even stronger cue-evoked activity, along with a modest decrease in reward-evoked activity among cue-excited neurons; however, there is also a marked increase in the proportion of reward-inhibited neurons, leading to a substantial reduction in the overall reward response. The observation that reward-evoked excitatory signaling peaks in adolescence might be consistent with adolescents’ elevated sensitivity to reward value; it also may reflect the finding that VTA dopamine neurons exhibit a much larger response to rewards, compared with adults, during Pavlovian (but not operant) conditioning (McCane et al., 2021).

Therefore, we would speculate that adolescent sign trackers, like adult sign trackers, use a dopaminergic RPE to transfer value from the reward to the cue over the course of adolescent training. The idea that acquisition of sign-tracking behavior involves an RPE-like learning process is supported by our finding that changes in behavior are best explained by the difference between cue-evoked and reward-evoked activity, rather than one or the other. In contrast to sign trackers, adolescent goal trackers and intermediates achieve the same transfer of neural activity and motivational value over the course of maturation into adulthood. Remarkably, this seems to occur with minimal additional training or experience of the cue, reward, or cue-reward contingency. This finding suggests that increased behavioral responding to cues in adulthood is not due to experience, per se, but to an altered balance between neural circuits promoting goal-directed action (e.g., goal tracking) and relatively inflexible, habit-like action based on the incentive salience of cues (e.g., sign tracking).

An important caveat is that this experiment cannot unequivocally distinguish between behavioral and neural changes resulting from development/maturation and those simply resulting from the passage of time. Indeed, anecdotal observations indicate that adult-trained subjects sometimes show increased sign tracking after a hiatus in training, possibly a phenomenon akin to the incubation of drug-seeking after abstinence (Chow et al., 2025). To disentangle these variables, future studies must compare behavior and neural cue responses before and after adolescent maturation with a similar time period of “abstinence” from Pavlovian conditioning in adulthood.

### Role of cue-evoked dopamine release

Consistent with previous findings in adult animals (Flagel et al., 2011; Lee et al., 2018), we found that NAc dopamine release in response to reward-associated cues was substantially higher in adolescent sign trackers compared with goal trackers. Unlike cue-evoked NAc neural activity, differences in dopamine release were present throughout all phases of training; indeed, we were surprised that sign trackers showed elevated cue-evoked dopamine during the very first training session. On the other hand, we have previously observed marked changes in reward-evoked activity over the first training session in adults (Gillis and Morrison, 2019) – which were present in sign trackers but not goal trackers – implying that dopamine-modulated neural circuits may undergo rapid adaptation during Pavlovian conditioning in certain individuals.

Similarly, unlike cue-evoked NAc activity, we found little change in cue-evoked dopamine release over the course of training among adolescent goal trackers, even as they matured into adults. However, this was consistent with the relatively small changes in behavior in this specific population of goal-tracker adolescents, which largely remained as goal trackers in adulthood. In contrast, subjects classified as intermediates showed a robust increase in cue-evoked dopamine release over the course of adolescent training, consistent with a behavioral trajectory that shifted from mainly goal tracking to a mix of sign tracking and goal tracking. We also noted that the magnitude of cue-evoked GRAB_DA_ signal was correlated with the strength of sign tracking, but not goal tracking, across subjects. This is in line with a number of previous findings indicating that the acquisition and expression of sign tracking, but not goal tracking, is dependent on both cue- and reward-evoked dopamine release in the NAc (Flagel et al., 2011; Saunders and Robinson, 2012; Chow et al., 2016; Iglesias et al., 2023; Herring et al., 2025).

Overall, the current study suggests that adolescents share with adults the same fundamental mechanisms of incentive salience attribution, which involves increasing cue-evoked dopamine and NAc activity and decreasing reward-evoked excitation in the NAc. However, our findings suggest that adolescents are less likely to engage this system, compared with adults, adding evidence to the hypothesis that adolescents are biased away from mesolimbic dopamine circuits (and possibly towards nigrostriatal circuits) in reward processing (McCane et al., 2021).

Additional research is needed to determine whether this is the result of developmental changes in the NAc – e.g., changes in dopamine receptor distribution and/or input from specific areas of mPFC – and/or changes in how/when the system for assigning incentive salience is gated by other processes, which might include hunger, social isolation, or other stressors (Anderson et al., 2013; DeAngeli et al., 2017).

